# Subjective salience ratings are a reliable proxy for physiological measures of arousal

**DOI:** 10.1101/2024.07.30.605866

**Authors:** Georgia E. Hadjis, Lauren Y. Atlas, Pedram Mouseli, Christine A. Sexton, Mary Pat McAndrews, Massieh Moayedi

## Abstract

Pain is an inherently salient multidimensional experience that signals potential bodily threats and promotes nocifensive behaviours. Any stimulus can be salient depending on its features and context. This poses a challenge in delineating pain-specific processes in the brain, rather than salience-driven activity. It is thus essential to salience match control (innocuous) stimuli and noxious stimuli, to remove salience effects, when aiming to delineate pain-specific mechanisms. Previous studies have salience-matched either through subjective salience ratings or the skin conductance response (SCR). The construct of salience is not intuitive, and thus matching through self-report poses challenges. SCR is used as a proxy measure that captures physiological arousal, which overcomes the nebulous construct of salience. However, SCR cannot be used to salience-match in real-time (i.e., during an experiment) and assumes an association between salience and physiological arousal elicited by painful and non-painful stimuli, but this has not been explicitly tested. To determine whether salience and physiological arousal are associated, thirty-five healthy adults experienced 30 heat pain and 30 non-painful electric stimuli of varying intensities. Stimuli were subjectively matched for salience and SCR was measured to each presentation. A linear mixed model found no differences in SCR between salience-matched heat and electric stimuli. A mediation analysis showed that salience fully mediated the relationship between stimulus intensity and SCR (proportion mediated=83%). In conclusion, salience and physiological arousal are associated, and subjective salience ratings are a suitable for salience-matching pain with non-painful stimuli. Future work can thus use subjective salience ratings to delineate pain-specific processes.

## Introduction

Pain is inherently salient [22], signaling potential or actual bodily threat [33]. Salience can be defined as the extent to which a stimulus differs from perceptual features of existing stimuli in the environment, making it stand out and grab our attention [41]. The salience of pain drives behaviour, first reorienting toward the stimulus [26], allowing the evaluation of the threat to mount an appropriate behavioural response. However, stimuli of any modality can be salient and elicit this cascade of events. Indeed, a series of studies showed no differences in brain activity elicited by salience-matched painful heat and non-painful electric stimuli [14,28,29]. Therefore, controlling for salience is essential to identify pain-specific processes.

Given the subjective nature of salience, it would be reasonable to rely on self-report ratings, as is the case with other subjective experiences (e.g., pain). However, cross-modal salience matching is not straightforward: a bright light may not grab the same level of attention as a noxious stimulus, given differences in the sensory qualia [2]. Furthermore, the concept of salience is nebulous, and thus not intuitive [1]. Thus, an objective measure to match physiological arousal as a proxy for salience across modalities could overcome this challenge.

The skin conductance response (SCR) is a phasic change in electrical conductivity of the skin elicited by an external stimulus [27]. SCR has previously been used to physiologically match salience between various levels of nociceptive and auditory modalities [13], demonstrating that stimulus intensity and intensity ratings are correlated, and increases in intensity ratings lead to increases in SCR magnitude. This study assumed that SCR magnitude approximates the extent of a stimulus’ salience. However, the association between physiological arousal and salience in response to noxious and innocuous somatosensory stimuli, and whether salience mediates the parametric relationship between stimulus intensity and SCR magnitude have yet to be formally tested.

Although desirable to have objective proxy measures of salience, salience-matching with these methods is determined after data collection during post-processing [4,9,13]—as opposed to matching in real-time using subjective salience ratings. This is particularly important in cases where post-processing reveals that salience was not adequately matched, resulting in data loss and data recollection to achieve adequate power [13] to parse out salience effects from effects of interest—i.e., pain-specific activity. However, if self-report salience ratings are tightly related to SCR magnitude across various stimulus intensities and modalities, this would allow real-time matching confirmation.

Here, we aimed to determine whether different intensities of subjectively salience-matched stimuli elicit similar levels of physiological arousal, as measured by SCR. Healthy participants experienced various intensities of two different somatosensory stimulus modalities—heat pain and non-painful electric shock—calibrated to elicit six target salience ratings, while SCRs were recorded. We hypothesized that SCR magnitude would increase as a function of self-report salience, and the SCR magnitudes for salience-matched heat pain and electric stimuli would not differ. We further hypothesized that increases in stimulus intensity would lead to increases in SCR, and that this would be mediated by salience ratings.

## Methods

### Participants

All study procedures were approved by the University of Toronto Human Research Ethics Board (Protocol #33465). Forty-seven individuals (24 female) aged 18-45 years were recruited from the University of Toronto environment for a single study visit. Sample size was determined using Bayes factor design analysis (BFDA; see *Salience and SCR AUC* below) [35]. Exclusion screening criteria were having current or past chronic pain, troublesome persistent or recurrent pain or pain-related discomfort in the last two months, clinically relevant levels of depression (> 21 on the Beck Depression Inventory; BDI), or reporting suicidal ideation (>0 on item 9 of the BDI). During the experiment, further exclusion criteria were: 1) perceiving electric stimuli as painful; 2) having high heat tolerance during calibration as we could not obtain a meaningful stimulus-response curve given ethical restrictions on thermal heat intensity (i.e., we cannot stimulate >51°C for an 8s stimulus); 3) reporting intensity ratings that were not consistent with increases with stimulus intensities during calibration; 4) being an SCR non-responder (see below); and 5) showing rapid SCR habituation defined as non-response to >2/5 stimuli, across any target salience levels.

### Study visit

Upon arrival to the study visit, participants provided informed e-consent and completed questionnaires. E-Consent and questionnaire data were collected and managed using REDCap electronic data capture tools hosted at the University of Toronto [10,11]. Questionnaires collected for exploratory analyses of individual differences included the TAHSN Demographic questionnaire, State Trait Anxiety Inventory (STAI) [36] and Pain Catastrophizing Scale (PCS) [38].

Our study was designed based on previous work by Horing and colleagues [13], with two key differences. First, we compared two somatosensory stimuli (noxious heat and innocuous electric stimuli, as opposed to auditory and heat pain) and our primary outcome measure was salience (which was not measured in the previous study). Second, while the goal was to obtain six unique salience ratings, the ratings were not grouped into different categorical levels in our analyses, as done by Horing et al. [13], but instead the actual ratings provided for each trial were used.

During the study visit, we first performed a familiarization procedure to determine the range of tolerated temperatures and currents. Next, to obtain six distinct salience ratings for the experiment, we performed a calibration procedure where the familiarization stimuli were rated for intensity, salience, and unpleasantness. We then performed a sigmoid regression using salience ratings to interpolate the stimulus temperatures and currents required to elicit the six target salience ratings during the experimental task.

During the experimental task, participants passively received 8-second heat or electric stimuli at six different intensities determined through interpolation. Participants’ SCR was recorded (see below) and they provided a salience rating after each stimulus.

### Familiarization

#### Thermal Stimuli

To familiarize participants with the various stimulus intensities used in the experiment, a familiarization procedure was performed for each modality. Thermal stimuli were administered throughout the experiment using a 9cm^2^ probe (TCS II T-11, QST.Lab, Strasbourg, France). The probe was held in a custom resin support fabricated by QST.Lab, and placed under the mid left calf. Temperatures 39°C, 41°C, 43°C, 45°C, 47°C, 48°C, 49°C were applied in ascending order at a ramp rate of 15°C/second for 8 seconds, and participants reported whether each stimulus was detected, painful, and tolerable (yes/no). If tolerance was reached before all temperatures were applied, the remaining temperatures were omitted. Participants were instructed of the IASP definition of pain, followed by “If the stimulus hurts or feels like it is or could potentially be damaging the skin, that would be considered painful.”

#### Electric Stimuli

Electric stimuli were administered using two surface electrodes placed 2cm apart along the sural nerve trunk on the right ankle. The threshold was determined by delivering single 200μs pulses (DS7A Digitimer, Welwyn Garden City, UK) starting at 1mA and steadily increasing the stimulus intensity by 1mA steps until the participant reported a light tapping sensation. Then, the current was lowered by 0.5mA until it was imperceptible. Next, current was increased by 0.25mA until another response was recorded, and then down and up by 0.05 until 5 positive responses were recorded. A train delay generator (DG2A, Digitimer) triggered the DS7A to deliver an 8-second train of electric stimuli (200 μs pulses) at 100Hz, and participants reported whether each stimulus was detected, painful, and tolerable (yes/no). The lowest intensity was at threshold followed by nine intensities that each increased by 0.5mA. If one of the stimuli was perceived as painful, the remaining currents were omitted. This was to ensure that currents used in the study were not perceived as painful.

### Rating Calibration

Next, a rating calibration procedure was performed, where participants received two rounds of nine stimuli per modality, first presented in ascending order, then again in a pseudorandom order. Stimulus parameters (duration, frequency, ramp rate) remained consistent with familiarization. Participants were asked to rate each stimulus along three rating scales: perceived intensity, perceived salience, and perceived unpleasantness. Intensity was defined as “the strength or magnitude of the stimulus.” Salience was defined as “the extent to which a stimulus is likely to grab and direct attention.” Unpleasantness was defined as “the extent to which a stimulus elicits a negative emotional experience.” Each scale ranged from 0-100 (intensity: 0 = not intense, 100 = most intense imaginable; salience: 0 = not salient, 100 = extremely salient, unpleasantness: 0 = not unpleasant, 100 = extremely unpleasant). Given the extremes of 100 on each scale, participants were instructed that providing a rating of 80/100 reflected the highest temperature or current they were willing to tolerate during the experiment. To distinguish the three rating scales from one another, a more detailed script was read to the participant (see Supplementary Methods).

#### Thermal Stimuli

The thermal probe and support were moved under the mid right calf. The maximum temperature was the highest tolerated temperature from familiarization, and the remaining eight temperatures each decreased by 1°C consecutively, for a total of nine temperatures. Consecutive stimuli were within 3°C of each other, and the highest three temperatures were not presented first, to mitigate bias in ratings [13].

#### Electric Stimuli

The same area as the familiarization procedure was stimulated, where the maximum current was the highest tolerated non-painful current from familiarization, and the remaining eight currents each consecutively decreased by 0.5mA, for a total of nine intensities. In cases where lower currents were perceived as painful, the stimulus intensity increments were adjusted to maintain an equal distance between stimuli with a minimum of 0.33.

Based on the ratings, a sigmoidal four parameter logistic (sigmoid) regression was performed for each participant to estimate stimulus intensities (temperatures and currents) needed to elicit a wide range of subjective salience ratings (25, 35, 45, 55, 65, 75) out of 100. In a separate sample, we demonstrated that this sigmoid regression model was a better predictor of salience ratings than a linear regression model for both stimulus modalities (see Supplementary Materials, Figure S1).

### Experimental Task

Participants experienced heat pain and non-painful electric stimuli at temperatures and currents calibrated to elicit six different salience ratings (Figure 1). Stimuli were presented in an alternating order of heat and electric, the target salience of which was organized in a pseudorandom order to repeat five times for each stimulus modality, for a total of 60 stimuli (30 electric, 30 heat). The stimuli were presented in a block design using PsychoPy v2023.2.3 on Windows [31] with 12 trials per block, for a total of 5 blocks. Given habituation to repeated stimuli, after the first two blocks, a second sigmoid regression determined the adjusted temperatures and currents to apply in the remaining three blocks. The pseudorandom order of target salience was determined using the same constraints as done previously [13]: two consecutive stimulus intensities had to elicit salience within three target rating values of each other; the block could not begin with intensities that elicited the highest two salience ratings. In each trial, a fixation cross on a laptop computer (IdeaPad 3 15IAU7, Lenovo, Beijing, China) would turn red to cue a stimulus event, a heat or electric stimulus would occur for 8 seconds, and then participants would provide ratings on an electronic visual analog scale to capture self-report ratings of intensity, salience, and unpleasantness, and answer whether the stimulus was painful (yes/no). Participants responded by moving a slider left or right using a mouse.

**Figure 1.**
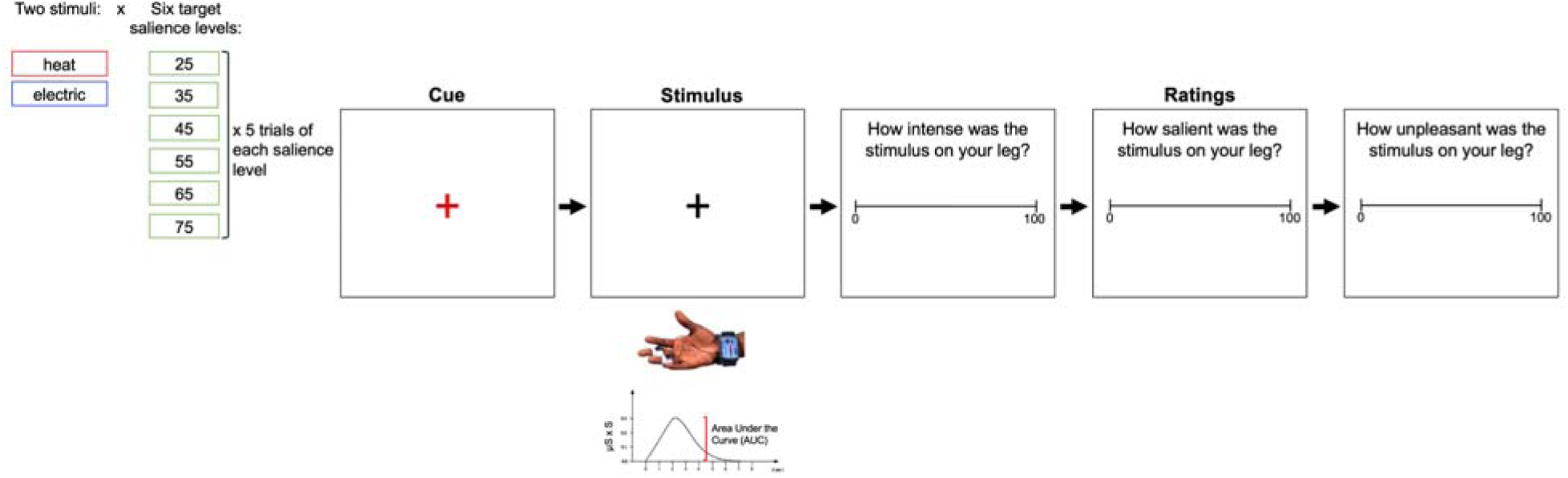
Experimental task design. As done previously, 8s stimuli were presented while their SCR was recorded, followed by a period where participants rated the perceived intensity, salience, and unpleasantness of the stimulus on a visual analog scale. Temperatures and currents that were projected to elicit six distinct salience ratings were applied 5 times for each stimulus modality (electric and heat), resulting in 60 total trials.

### SCR Recording Setup

During the experimental task, SCRs were recorded using the MP160 Data Acquisition System and Bionomadix wireless PPG/EDA amplifier (BIOPAC Systems Inc., Goleta, CA, USA). Disposable electrodermal activity (EDA: EL507) isotonic gel electrodes were applied with isotonic electrode gel to the index and middle fingertips of the left hand. A wireless Bionomadix transmitter was worn around the wrist and connected to the electrodes using EDA electrode leads. The transmitter sent a signal to the amplifier which was recorded at a sampling rate of 62.5 Hz on a separate laptop computer (XPS 15-9570, Dell Technologies, Round Rock, TX, USA) on BIOPAC’s Acqknowledge v.5.0 software. Once connected, we performed a final screen before the experiment to ensure that participants were SCR responders. To do so, participants were instructed to take a deep breath, which generates a visible SCR. Two of the twelve excluded participants who consented to procedures did not demonstrate a visible SCR and were SCR non-responders.

The PsychoPy task triggered the devices (TCS II or DS7A) to deliver the specified stimulus modality and intensity for each trial by sending a TTL pulse via a Multifunction Input/Output Device (NI USB-6343, National Instruments Corp., Austin, TX, USA). Given the manual control of stimulus intensity of the DS7A, the experimenter would set the next stimulus current to be applied during the rating screen of the previous stimulus. The stimulation devices each sent a digital TTL pulse to the MP160 unit such that each stimulus modality had its own recording channel in Acqknowledge, marking the onset and offset of each stimulus.

### SCR Analysis

SCR data was analyzed with the Acqknowledge v5.0 software. First, we performed a low-pass filter at 1Hz to remove noise and high frequency signals in the waveforms. We then visually inspected data for artifacts in the recorded waveform during the stimulus event, such as sudden finger twitch and hand movements, and connected waveform endpoints to smooth the signal. The stimulus event was used to estimate event-related SCRs, which were defined as the SCRs occurring 0.5-14 seconds after stimulus onset. Event-related SCRs with amplitudes below 0.02μs and those with no SCR responses were considered non-responses, but were kept in the analysis [24]. We measured the area under the curve (AUC) [24] of event-related SCR amplitudes for every trial and organized them across stimulus modality and target salience rating per participant. In some cases, SCRs within the time window were not captured by the software, and we commissioned a script from BIOPAC to address this issue.

### Stimulus Intensity and Salience Ratings

To determine whether stimulus intensity (temperature and current applied on each trial) elicited a parametric increase in self-report salience, we performed a linear mixed model with salience rating as the dependent variable, stimulus intensity as a fixed effect, and subject as a random effect. Separate linear mixed models were run for temperature and current as stimulus intensity. Stimulus intensity was mean-centered across all trials of each modality to ensure meaningful interpretations of the intercept. Significance was set at p < 0.05.

### Rating Scale Comparisons

Given the relatedness of the rating scales used in the experiment (intensity, salience, and unpleasantness), it is important to ensure participants were able to conceptually distinguish them. One way to do so is to determine how similar ratings are within participants and whether there are statistical differences in these self-report values. Thus, we performed a Spearman correlation and Wilcoxon signed-rank test to assess similarity and differences, respectively. We used nonparametric tests given that the data were not normally distributed (Shapiro Wilk test, p < 0.05). Significance was set a p < 0.05.

### Salience and SCR AUC

*Sample size*: Sample size was determined with design BFDA [35] using the BFDA package in RStudio [34], where interim analyses are performed at pre-determined proportions of a final sample size (25%, 50%, and 75%), and data collection ends when a Bayes factor (BF) with moderate to strong evidence is obtained supporting either the null or the alternative hypothesis. Based on this approach, 66 subjects were projected to obtain a moderate to strong BF with 80% power. To obtain a BF, we performed a 2 (stimulus modality: heat vs electric) x 6 (salience: 25, 35, 45, 55, 65, 75) Bayesian repeated measures ANOVA (JASP Team 2024, Version 0.19[Intel]) that assessed a null model against several other models, all of which included the random effect of subject:

(1) the null model only included subject as a random effect and no effect of or relationship between salience or stimulus modality on SCR AUC;
(2) a stimulus modality model, where stimulus modality alone affects SCR AUC;
(3) a salience model, where salience alone affects SCR AUC;
(4) a stimulus modality and salience model, where each factor independently affects SCR AUC; and
(5) a stimulus modality and salience interaction model, where each factor and their interaction affects SCR AUC.

A BF_10_ value was calculated for each model compared to the null model, where values greater than 1 indicate support for the alternative model, and values lower than 1 indicate support for the null model. We then performed an analysis of effects which averages across the models to determine the evidence for each effect using a BF inclusion value. The Bayesian ANOVA was run using salience ratings only, as that was the scale of interest that determined the sample size.

To determine whether salience ratings and SCR AUC were related, and whether there were differences in this relationship between modalities, we fit a random slope and intercept linear mixed model using the lmerTest package in RStudio (RStudio, Boston MA, USA) [17]. Specifically, the dependent variable was AUCs and independent variables were stimulus modality and salience rating. Two further linear mixed models were performed, one for intensity ratings and one for unpleasantness ratings, with the same predictors. The fixed effects for each model were stimulus modality and rating, and participant was set as a random effect. Significance was set at p < 0.05.

### Mediation Analysis

Upon observing a relationship between stimulus intensity and salience, and a relationship between salience and SCR AUC independent of stimulus modality, we performed a mediation analysis to assess whether there was an association between stimulus intensity and SCR AUC, and the extent to which it was mediated by salience. The temperature and current applied on respective heat and electric trials was z-scored to transform the data between modalities into comparable units. The independent variable, *X*, was the z-scored stimulus intensity (temperature/current) applied on a given trial, the mediator, *M*, was the subjective salience rating in that trial, and the dependent variable, *Y*, was the SCR AUCs elicited by the stimulus applied in that trial. First, three linear regressions were performed to ensure that *X* predicts *Y* (Path *c*) and *X* predicts *M* (Path *a*)*—*prerequisites for a mediation analysis. Next, the indirect mediation effect of *M* on *Y* while controlling for X (Path *b*) was assessed, which also allowed us to compute the indirect effect (Path a*b, mediation) and the direct effect (c’, effect of X on Y while controlling for the mediator). Statistical significance of the mediation was computed using bootstrapping with 1000 permutations with the mediate function in the RStudio mediation package [40]. The same procedures were performed for perceived intensity and perceived unpleasantness ratings, separately. Significance was set at p < 0.05.

## Results

### Participants

We recruited 47 participants, of which 12 were excluded based on the following criteria:

(1) participants perceived electric stimuli as painful (n = 1);
(2) given ethical restrictions on thermal heat intensity (i.e., we cannot stimulate >50°C for an 8s stimulus), participants who had high heat tolerance during calibration were excluded as we could not obtain a meaningful stimulus-response curve (n = 2);
(3) intensity ratings were not consistent with increases with stimulus intensities during calibration (n = 1);
(4) participants were SCR non-responders (n = 2);
(5) participants showed rapid SCR habituation defined as non-response to >2/5 stimuli, across any of target salience levels (n = 4); and
(6) participants’ data were analyzed to finalize the analysis pipeline (n=2).

At 35 participants with full datasets, we performed an interim analysis. The Bayesian repeated measures ANOVA revealed that the factors included in the model met sufficient evidence criteria, specifically the salience model (BF_10_ = 129.8) and the salience and modality model (BF_10_ = 18.6) (see Table S1). This was further supported by an analysis of effects which showed that only the main effect of salience had evidence for the alternative hypothesis (BF_incl_ = 87.42), whereas the main effect of modality and interaction between salience and modality showed evidence for the null hypothesis (BF_incl_ < 0.3; see Table S2). The final sample of 35 participants comprised 19 females, 16 males, mean age ± SD = 25 ± 4.65 years, range: 18-38 years. Demographic data are provided in Figure 2 and Table 1. Questionnaire data are provided in Table 2.

**Figure 2.**
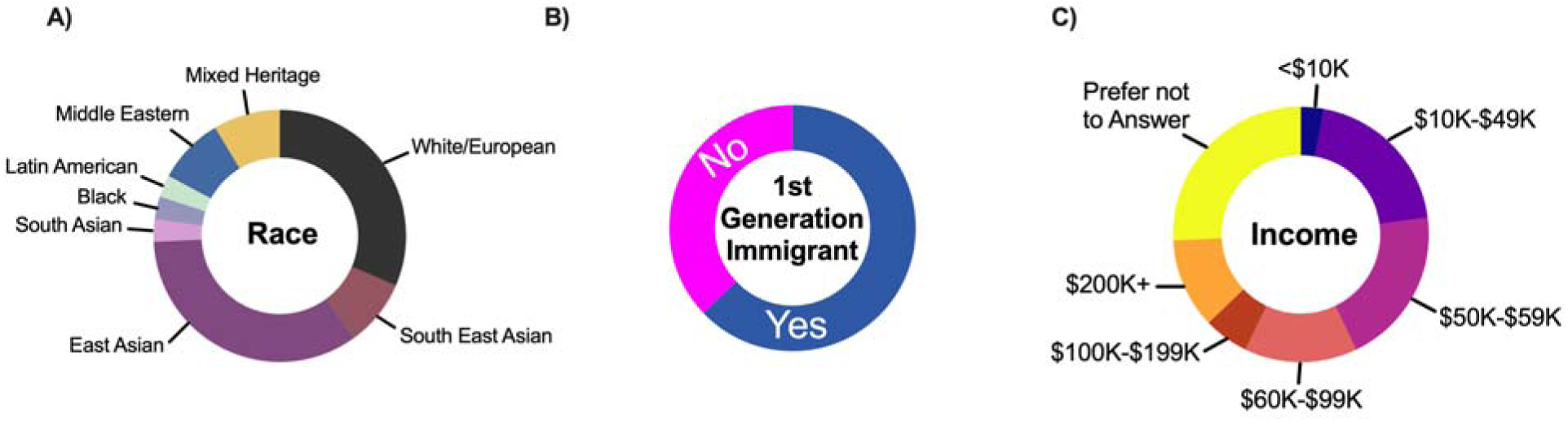
Demographic data. (A) Distribution of racial groups in the sample. (B) First generation immigrant status. (C) Distribution of income supporting 2.4 individuals per household.

**Table 1.**
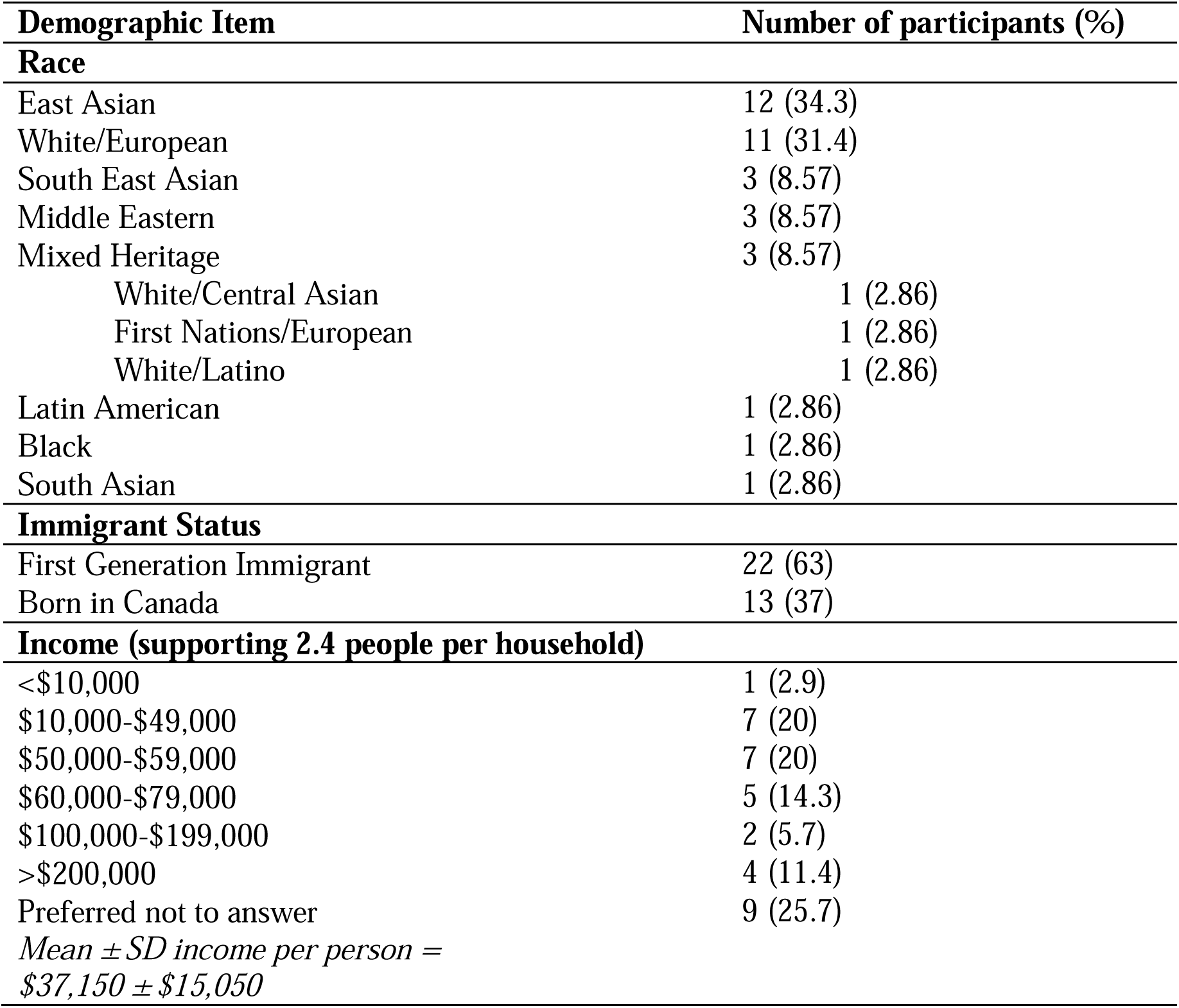
Participant Demographics.

| **Demographic Item** | **Number of participants (%)** |
| --- | --- |
| **Race** |  |
| East Asian | 12 (34.3) |
| White/European | 11 (31.4) |
| South East Asian | 3 (8.57) |
| Middle Eastern | 3 (8.57) |
| Mixed Heritage | 3 (8.57) |
| White/Central Asian | 1 (2.86) |
| First Nations/European | 1 (2.86) |
| White/Latino | 1 (2.86) |
| Latin American | 1 (2.86) |
| Black | 1 (2.86) |
| South Asian | 1 (2.86) |
| **Immigrant Status** |  |
| First Generation Immigrant | 22 (63) |
| Born in Canada | 13 (37) |
| **Income (supporting 2.4 people per household)** |  |
| <\$10,000 | 1 (2.9) |
| \$10,000-\$49,000 | 7 (20) |
| \$50,000-\$59,000 | 7 (20) |
| \$60,000-\$79,000 | 5 (14.3) |
| \$100,000-\$199,000 | 2 (5.7) |
| >\$200,000 | 4 (11.4) |
| Preferred not to answer | 9 (25.7) |
| *Mean ± SD income per person =*<br>*\$37,150 ± \$15,050* | |

**Table 2.**
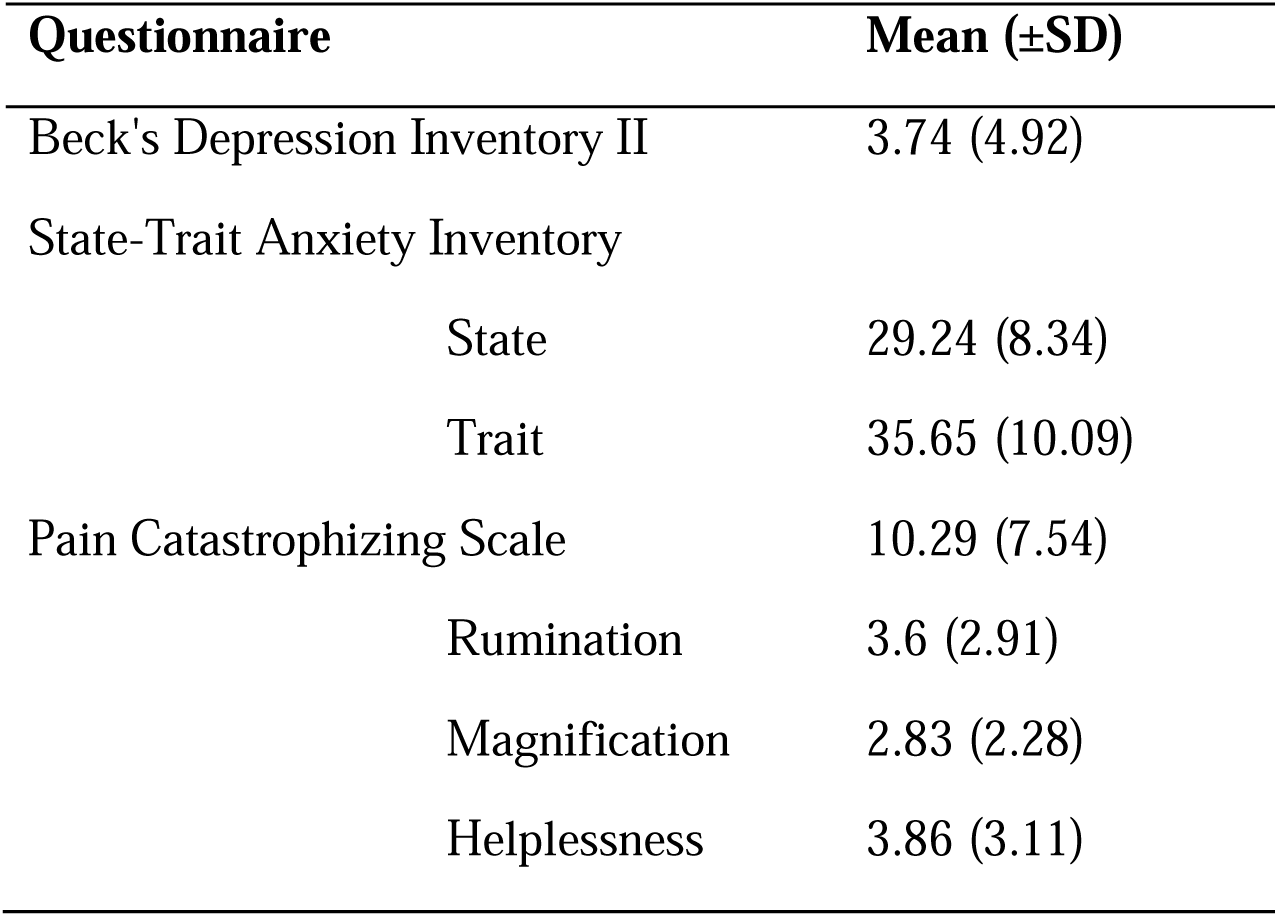
Group Means of Self-Report Questionnaires.

### Salience ratings increase as a function of stimulus intensity

As shown in Figure 3, linear mixed models revealed that as temperature and current increase, self-report salience ratings increase (electric: β = 23.92, t = 12.14, p = 1.34e-13; heat: β = 15.17, t

**Figure 3.**
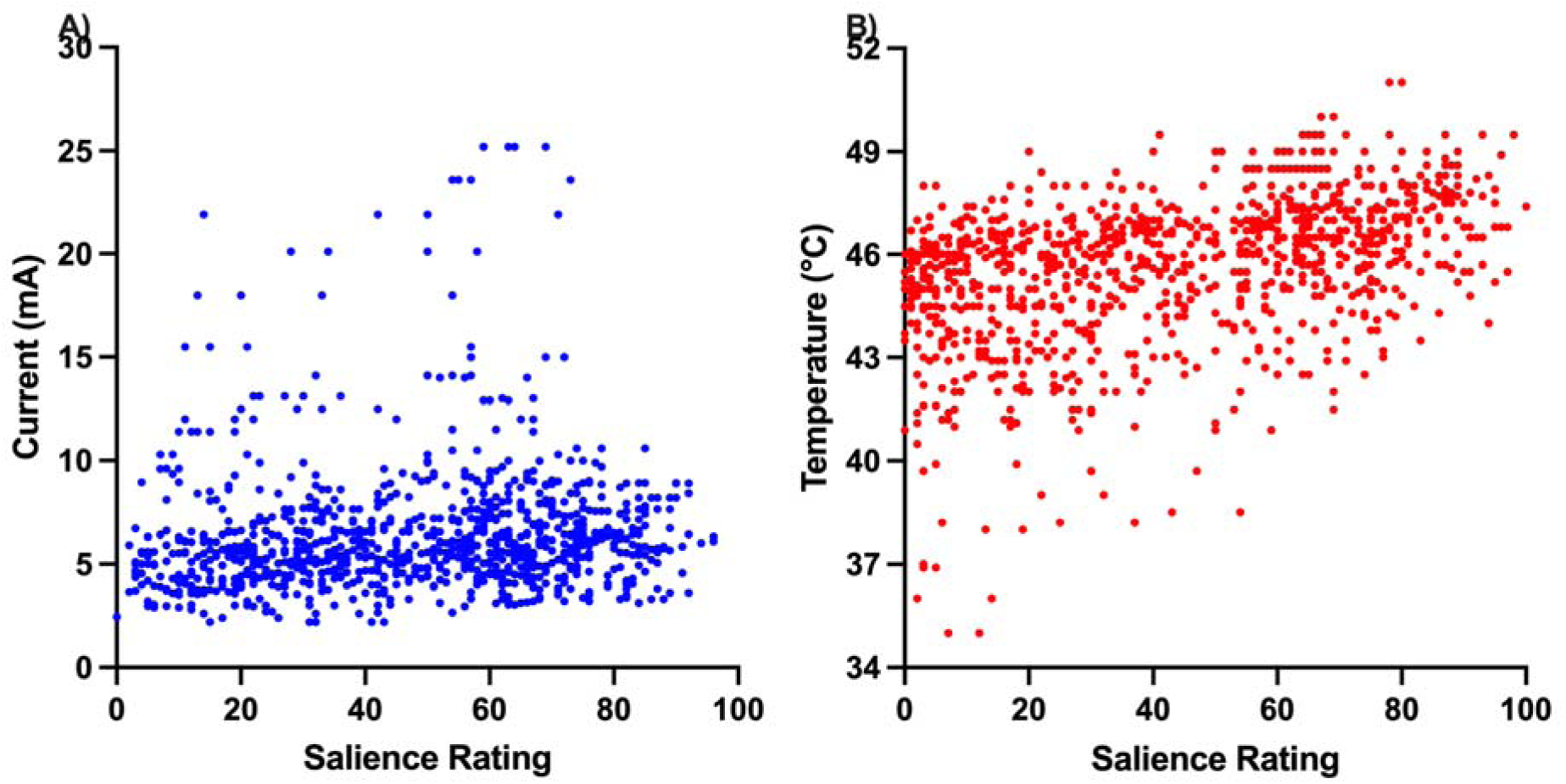
Relationships between stimuli intensity and salience ratings. Stimulus intensities (A) electric current (mA) and (B) temperature (°C) and corresponding salience ratings for each trial for all participants.

= 10.8, p = 1.41 e-11).

### Rating Scales are correlated but their medians differ

Rating scales were all significantly correlated (Figure 4): salience vs intensity: *rho* = 0.93, p < 0.001; salience vs unpleasantness: *rho* = 0.79, p < 0.001, intensity vs unpleasantness: *rho* = 0.81, p < 0.001. Wilcoxon signed-rank tests revealed that the median for salience ratings was greater than the median of intensity ratings (Z = 19.20, p < 0.001), median salience ratings were greater than median unpleasantness ratings (Z = 32.42, p < 0.001), and median intensity ratings were greater than median unpleasantness ratings (Z = 24.35, p < 0.001).

**Figure 4.**
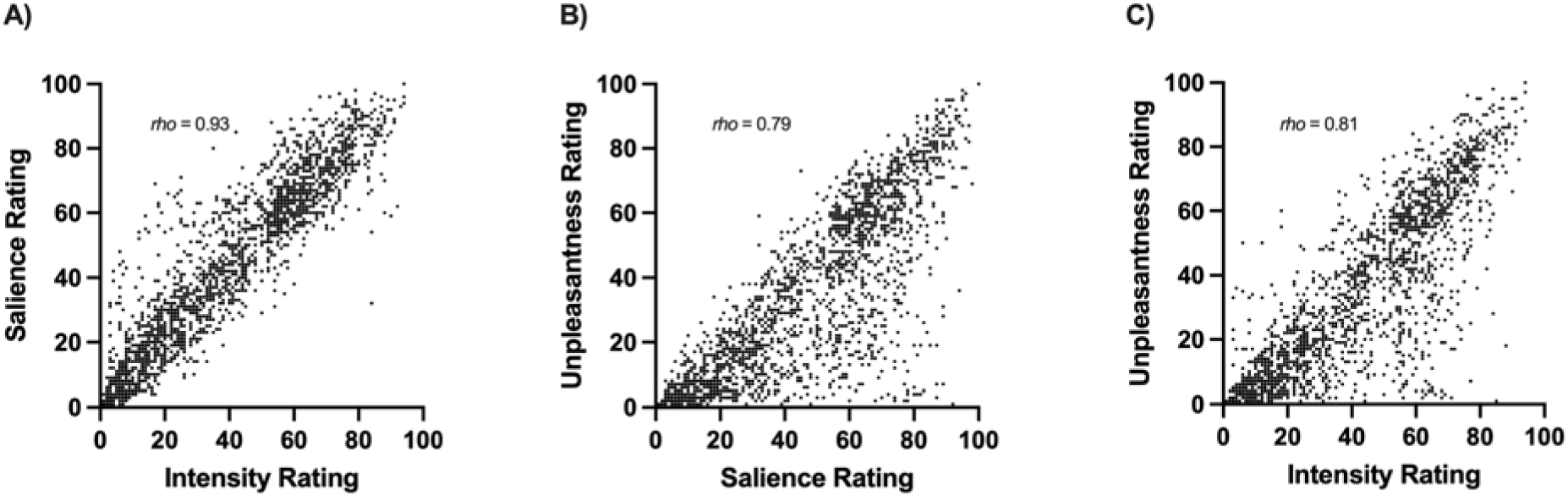
Relationships between ratings scales. Correlations between (A) intensity and salience, (B) unpleasantness and salience, and (C) intensity and unpleasantness ratings on each experiment trial per participant.

### SCR is associated with salience ratings

A linear mixed model comparing the relationship between salience and stimulus across modalities (i.e., heat vs. electric) revealed a significant main effect of salience (β = 0.023, t = 4.45, p = 8.55e^-05^, η^2^ = 0.41), but no main effect of stimulus modality (β = 0.0096, t = 0.049, p = 0.96) or stimulus-by-salience interaction (β = 0.0048, t = 0.74, p = 0.47). These results demonstrate that as self-reported salience increases, there is a corresponding increase in SCR, a measure of physiological arousal, indicating that salience and SCR are associated (Figure 5). In addition, these results suggest that subjectively salience-matched electric and heat stimuli can in fact be matched physiologically, as the SCR AUCs between stimulus modalities do not differ as salience increases. Results for intensity and unpleasantness ratings, which were also associated with SCR, can be found in the Supplementary Materials.

**Figure 5.**
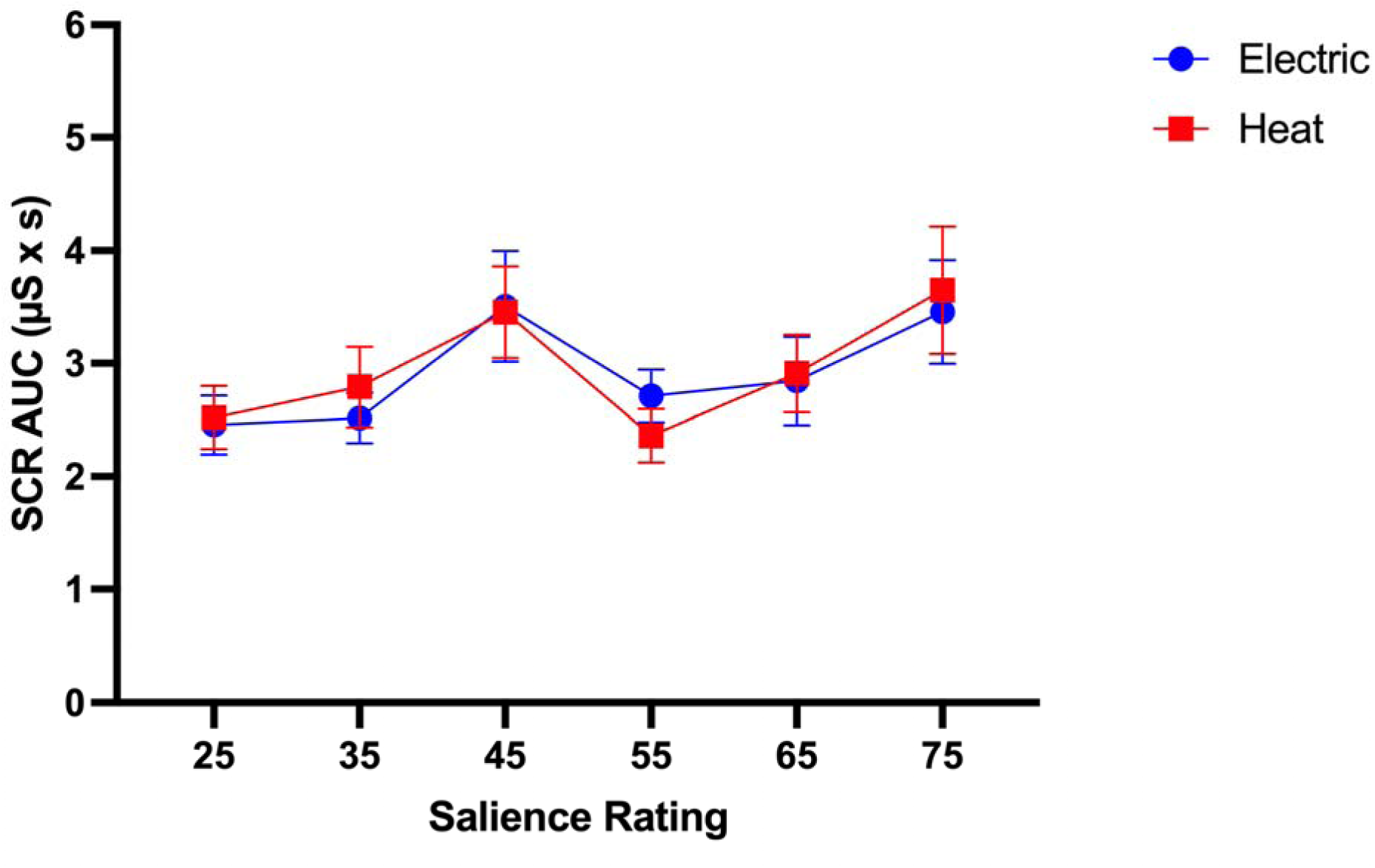
Area under the curve (AUC) of the skin conductance in response to heat (red) and electric (blue) stimuli across the six self-report salience ratings. Group mean ± SEM AUC in μSiemens-by-second is plotted for trials meant to elicit each target salience rating.

### Salience Mediates the Relationship between Stimulus Intensity and SCR

The mediation analysis (Figure 6) revealed that stimulus intensity had a significant effect on salience (*a* = 16.41, p < 2e^-16^), and the total effect of stimulus intensity on SCRs was significant (*c* = 0.44, p < 0.0001). However, when accounting for salience as a mediator, salience had a significant effect on SCRs (*b* = 0.022, p < 2e^-16^), but stimulus intensity no longer predicted SCR (*c’* = 0.073, p = 0.38). This was confirmed by causal mediation analysis, where we found that salience completely mediated the relationship between stimulus intensity and SCR (*ab* = 0.36, p < 2e^-16^; proportion mediated = 83%), as there was no direct effect of stimulus intensity on SCRs.

**Figure 6.**
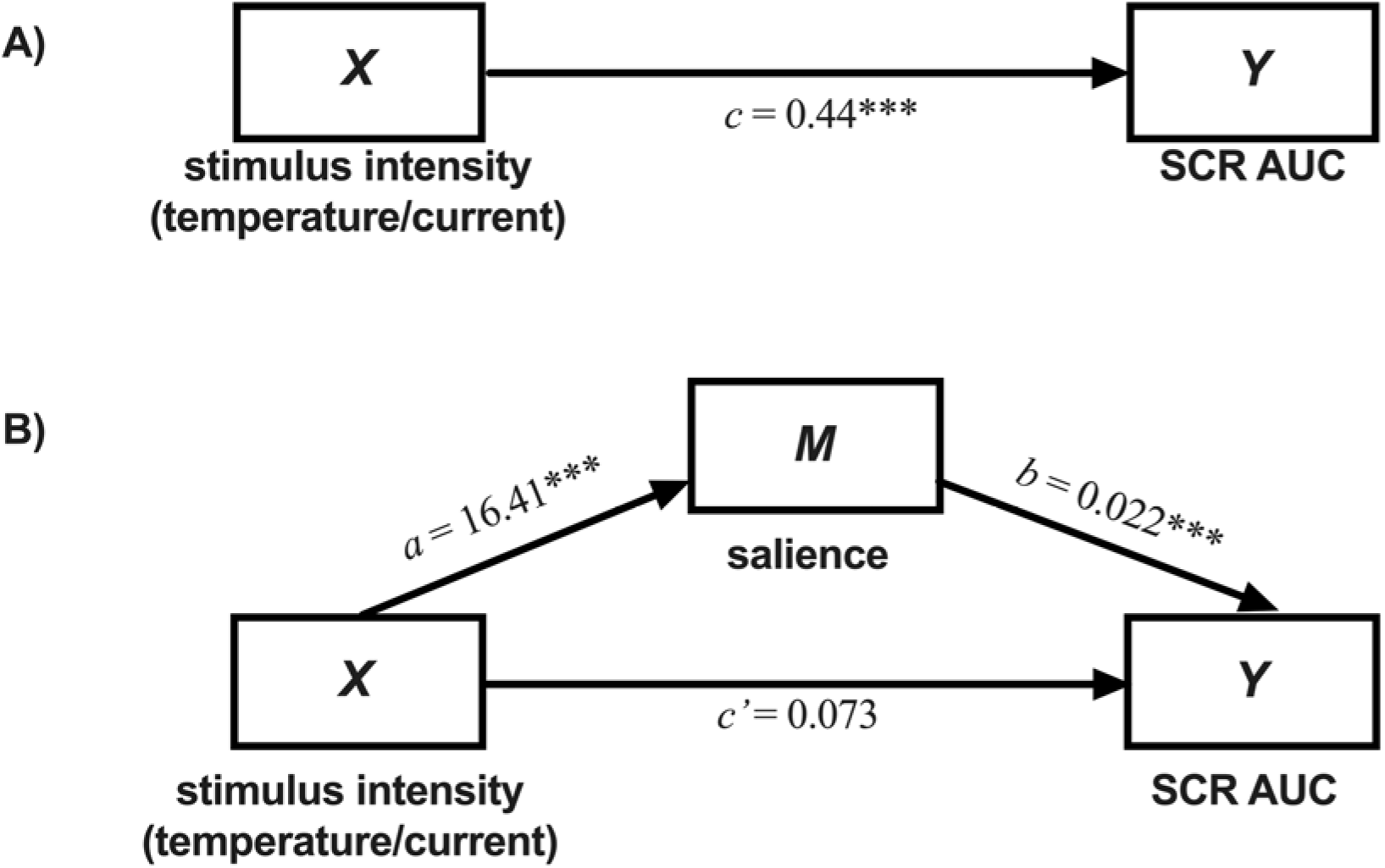
Mediation results where salience mediated the relationship between stimulus intensity and SCR AUC. (A) The total effect of stimulus intensity (*X*) on SCR AUC (*Y*). B) Regression coefficients between each factor, where *a* and *b* represent the indirect effect (mediation), and *c’* represents the direct effect of stimulus intensity on SCR AUC accounting for and SCR AUC, respectively.

These findings are evidence that salience is directly associated with and contributes to the increase in physiological arousal that occurs as stimulus intensity increases. The reverse analysis with SCR as the mediator and salience as the outcome did not reveal mediation (*c’* = 15.56, p < 2e^-16^, proportion mediated = 4.9%). Mediation results for intensity and unpleasantness ratings, which also showed complete mediation, are reported in the Supplementary Materials (Figure S2).

## Discussion

This study aimed to determine whether subjective salience ratings are associated with a physiological measure of arousal, SCR, in response to somatosensory stimuli of different modalities and intensities. We found that as stimulus intensity increased, salience ratings increased, and increases in salience ratings were associated with increases in SCR. These increases in SCR were not different between modalities, allowing for salience-matching. A causal mediation analysis revealed that the relationship between stimulus intensity and SCR was completely mediated by salience. These findings indicate that self-report salience ratings are a suitable metric to track physiological arousal. Self-report can thus be used to perform reliable, real-time salience-matching in studies where matching is required across modalities.

When investigating pain-specific brain processes, and their association with behaviour, researchers face the challenge of parsing pain-specific activity from the salience content of the painful stimulus [6,20–22]. Salience alone is sufficient to drive behaviour—reorienting toward the stimulus [3,26], evaluating of the potential threat, and mounting an appropriate behavioural response. To identify pain-specific processes, controlling for salience is essential. One approach to do so is to salience-match an innocuous stimulus and a noxious stimulus. This has previously been done in a series of studies that showed no differences in brain activity using mass-univariate statistics in response to salience-matched heat and electric stimuli [14,28,29]. In these studies, salience was loosely defined to participants as “the ability of the stimulus to capture attention.” However, there are significant limitations to this approach: the nebulous definition of salience, the unfamiliarity of the term for the average person [1], and the lack of validation of self-report salience ratings. It is not clear how an individual could assess the extent to which a stimulus grabs their attention on a scale from 0 (“not salient”) to 10 (“extremely salient”), without training or comparators.

One approach that has been utilized to overcome the limitation of subjective salience-matching is to record objective, physiological arousal in response to pain [25] and innocuous stimuli using the skin conductance response (SCR) [13]. In these studies, the SCR is considered a proxy measure of arousal [27]. In pain research, it is used as a measure of physiological arousal from noxious stimuli [5,7,8,37,39], and as an indicator of pain in non-communicative populations [30,37]. Indeed, Geuter et al [8] showed that the temporal profile of SCR can be used to predict trial-by-trial pain ratings, but again, this does not allow for the isolation of pain-specific activity—and, if anything, it shows a physiological signature of pain ratings, but does not explicitly test the association between SCR and salience.

A significant limitation to using SCRs as a measure of salience-matching is this cannot be performed during the experiment, and requires post-processing and analysis to determine whether matching was successful. This can lead to data or participants being excluded, which can be costly as well as resource- and time-intensive. Here we address this limitation by obtaining both SCRs and self-reported salience to determine whether subjective salience tracks with SCR, as a measure of physiological arousal. In addition, collecting self-report ratings allowed us to perform real-time salience-matching during the experiment, circumventing the data loss that can occur when salience-matching post-hoc using SCRs only. Our methods follow the procedures conducted by Horing et al. [13], with a few key differences: we compared the salience of pain to that of an innocuous electric stimulus, rather than an auditory stimulus, and the self-report rating of interest was perceived salience, rather than perceived intensity of heat and unpleasantness of the control stimulus.

To perform subjective salience-matching properly, it is essential to operationalize salience with a clear, understandable definition at the beginning of the experiment, as well as providing contextual examples. Here, participants were provided an explanation with analogies to help understand how to interpret salience. We operationalized salience as “the extent to which the stimulus grabs and directs your attention” (see Supplementary Materials for the precise script used). In other words, as they felt the stimulus on their leg, they were asked to think about how much of their attention they allocated to that sensation. We also differentiated salience from perceived intensity and unpleasantness by providing a simple thought exercise: participants were instructed to imagine that they are studying intently in a quiet library and suddenly experience a light tap on their shoulder from a friend trying to get their attention. This example highlights to participants that a stimulus can be salient without being intense or unpleasant. No difference in SCRs between stimuli across each rating scale indicates that the scales are related, which is expected in healthy adults. While we show that intensity and salience ratings are highly correlated, they do not assess the same sensory quality, and correlating intensity ratings with physiological arousal does not answer whether salience and arousal are associated [14,15,20,28,29]. In fact, we found that the medians of the three rating scales were significantly different from one another, suggesting that participants distinguished the scales while also understanding the ways in which they are conceptually correlated (i.e., an unpleasant stimulus is likely to be intense and salient). Our mediation results show that each rating scale completely mediated the relationship between stimulus intensity administered and SCRs, which provides novel and reproduced evidence that perceived salience and intensity/unpleasantness are associated with physiological arousal, respectively (Figure S2).

Much like any subjective experience, the salience of a stimulus is dynamic, and ratings are often inconsistent over time [4,12,16,23,32]. Therefore, it is important to consider the frequency of sampling of salience ratings when aiming to salience-match across stimuli. Asking for ratings too often can lead to the participant paying too much attention to the stimulus, and thus amplifying the experience. Conversely, sampling too sparsely can miss changes in salience, and adversely affect matching. Therefore, salience-matching may not be maintained over an experiment, and taking ratings only at the beginning or end of the experiment will miss these dynamic changes in the sensory experience. One way to overcome this is to parse an experiment into multiple blocks, and assess the salience of stimuli after each block. This provides an opportunity to adjust stimulus intensity based on ratings provided thus far by using stimulus response curves to maintain the desired salience rating.

The present study is unique in that we compared the objective SCR to increasing levels of salience-matched noxious and innocuous stimuli to determine whether subjective and physiological arousal are in fact associated. We found no differences in physiological arousal in response to heat and electric stimuli as the salience increased, represented by the AUC of the SCR. These findings are in line with previous work showing no difference in SCR between heat pain and auditory stimuli [13], and SCR increasing as a function of numeric rating scale reports of postoperative pain [18,19]. Importantly, we show that salience completely mediated the increase in physiological arousal that occurs because of increasing stimulus intensity, providing further evidence of the strong association between SCRs and salience. Here, we demonstrate that salience is associated with physiological arousal, and when instructed properly, self-reported salience ratings can be used as a proxy of physiological arousal to investigate pain-specific processes.

A common limitation of using physiological arousal as an outcome measure is the habituation to novelty, and while a repeated block design is essential for sufficient within-subject power, it becomes predictable. SCRs as one such measure of physiological arousal are known to habituate quickly to external stimuli and become smaller and less frequent over time. In addition, SCRs are greater when individuals are particularly nervous or excited. As such, when participants become more comfortable with study procedures and the repetition of stimulus trials, arousal decreases, and SCR decreases as a result. Additional measures of arousal such as pupil dilation may provide clarity on the physiological responses to stimuli over time.

In sum, we show that salience and physiological arousal are associated. Subjectively salience-matching pain with innocuous stimuli, when instructed properly, can be used to control for salience while investigating pain-specific processes, as it provides a real-time metric that can be adjusted to reflect the dynamic experience of pain.

## Acknowledgements

M Moayedi is supported by a University of Toronto Centre for the Study of Pain — Pain Scientist Award, a Canada Research Chair (Tier 2) in Pain Neuroimaging, and the Bertha Rosenstadt Endowment Fund at the Faculty of Dentistry, University of Toronto. The authors have no conflicts to report. This study was funded through Moayedi’s discretionary funds. LY Atlas is funded through the Intramural Research Program of the National Center for Complementary and Integrative Health (ZIA AT000030). GE Hadjis is supported by the Ontario Graduate Scholarship, Queen Elizabeth II Graduate Scholarship in Science and Technology, Lupina Foundation Health & Society Bursary, Canadian Institutes of Health Research Canada Graduate Scholarship-Master’s, the Harron Fund at the Faculty of Dentistry at the University of Toronto, and the Pain Scientist Scholarship from the University of Toronto Centre for the Study of Pain.

We thank Dr. Tim V. Salomons for help in developing the script to distinguish between stimuli rating scales and Ms. Reem Mustafa for data collection for a previous version of this manuscript.

## Data availability statement

Data are available upon reasonable request.

## Notes

### Competing Interest Statement

The authors have declared no competing interest.

### Summary of Updates

1. Revised the overall objective/purpose of the manuscript for consistency 2. Methods updated for clarity and to enable reproducibility. 3. Collected new data, with sample size based on a Bayes Factor Design Analysis 4. Revised the rating scale a more traditional rating scale. Our rating scale considers a stimulus at 80/100 as tolerance, and we did not deliver stimuli that would elicit ratings greater than 80/100. 5. We added a demographics questionnaire and now report data on sex, age, race, income, and immigrant status. 6. We recruited an independent sample of participants to evaluate whether a linear or sigmoid regression would better interpolate target salience ratings (25, 35, 45, 55, 65, 75) for the experimental task. Sigmoid regression better predicted the stimulus intensities that elicited the target salience ratings, and this sigmoid was used in the main study as a result. 7. Addition of a control mediation analysis of the opposite pathway. 8. Author list updated

