## Supplementary for "Subjective salience ratings are a reliable proxy for physiological measures of arousal"

^1^Centre for Multimodal Sensorimotor and Pain Research, Faculty of Dentistry, University of Toronto, ON, Canada M5G 1G6;  ^2^National Center for Complementary and Integrative Health, National Institutes of Health, Bethesda, MD, USA^; 3^National Institute of Mental Health, National Institutes of Health, Bethesda, MD, USA, ^4^National Institute on Drug Abuse, National Institutes of Health, Baltimore, MD, USA; and ^5^Division of Clinical and Computational Neuroscience, Krembil Brain Institute, Toronto Western Hospital, University Health Network, Toronto, ON, Canada, M5T 2S8; ^6^University of Toronto Centre for the Study of Pain, Faculty of Dentistry, University of Toronto, Toronto, ON, Canada M5G 1G6

**Running title (45 characters including space):** Salience ratings and SCR

Data Availability: Anonymized data are available upon request

§Please address all correspondence to: Massieh Moayedi

Centre for Multimodal Sensorimotor and Pain Research

Faculty of Dentistry, University of Toronto

123 Edward Street, Room 501B

Toronto, ON

Canada M5G 1E2

Bluesky: massih.bsky.social

Website: [www.painresearchcentre.org](http://www.painresearchcentre.org/)

**Supplemental Methods**

We use the following script to describe salience to participants, and to help them understand the difference between intensity, unpleasantness and salience.

Salience, unpleasantness, and intensity are related to each other. It is possible for something to be salient, unpleasant, and intensely painful all at the same time. Touching a hot stove with your finger will capture your attention and is very unpleasant. It is unpleasant, in part, because of the feeling of pain associated with the burn injury to your finger. Essentially, salience is how much of your attention is drawn towards the stimulus.

On the other hand, salience, unpleasantness, and pain don’t necessarily occur together. It is possible for something to be salient but not unpleasant or painful. For example, if you are studying for an upcoming test in a library and a friend taps you on the shoulder, it will grab your attention, but it does not (usually) cause a negative emotional response. It also doesn’t feel like the tapping is going to cause you physical harm or cause injury. Thus, the tap is salient, but not unpleasant or painful.

It is also possible for something to be unpleasant but not painful. Hearing a tragic news story might make you feel sad, but it doesn’t feel like any part of your body is being damaged or suggests you need to protect yourself from physical injury. Thus, the news story is unpleasant but doesn’t cause you pain.

The distinction between intensity and unpleasantness is clear if you think of listening to a sound, such as a radio. As the volume of the sound increases, I can ask you how loud it sounds or how unpleasant it is to hear it. The intensity of pain is like loudness; the unpleasantness of pain depends not only on intensity but also on other factors which may affect you. Some pain sensations may be equally intense and unpleasant, we would like you to judge the two aspects independently.

Is the distinction between salience, unpleasantness, intensity, and pain clear? Do you have any questions?

You are about to experience a series of stimuli. These stimuli might capture your attention, but will not necessarily be unpleasant or painful. If the stimuli do not bother you, do not rate them as unpleasant. If the stimuli don’t physically hurt, do not rate them as painful.

**Linear vs Nonlinear models of salience ratings**

To assess whether linear or four-parameter logistic (4PL) curve sigmoid regression better captured the relationship between salience ratings and stimulus intensity, the same familiarization as the main study and five rounds of the rating calibration procedures were performed in a separate sample. Temperatures and currents interpolated by linear and sigmoid models were then applied three times each in a pseudorandom order, and the corresponding salience ratings were fit using linear and sigmoid models. To ensure robust model evaluation and prevent overfitting, a leave-one-out cross-validation (LOOCV) scheme was employed. At each iteration, one data point was excluded for testing while the models were fitted to the remaining data points. Predictions for the left-out samples were collected to compute the mean squared error (MSE) for each model and participant. The comparison of MSE values across models allowed for an objective assessment of model fit.

**Supplemental Results**

**Linear vs Nonlinear Regression**

In a sample of 8 healthy adults (4 female, mean ± SD = 22.5 ± 2.56), the LOOCV scheme revealed that the sigmoid regression better predicts and fits salience ratings (Figure S1) for both electric (average linear MSE = 189.4; average nonlinear MSE = 167.3) and heat stimuli (average linear MSE = 312.5; average nonlinear MSE = 180.1).

**Linear Mixed Model**

There was a main effect of intensity (*β* = 0.026, t = 4.19, p = 0.00014, η^2^ = 0.44) but no main effect of stimulus (*β* = 0.0095, t = 0.048, p = 0.96) or stimulus-by-intensity interaction (*β* = 0.0044, t = 1.05, p = 0.30). Similarly, there was a main effect of unpleasantness (*β* = 0.027, t = 3.76, p = 0.00053, η^2^ = 0.35) and no main effect of stimulus (*β* = -0.11, t = -0.69, p = 0.49) or stimulus-by-unpleasantness interaction (*β* = 0.0035, t = 0.89, p = 0.38). As with the results of the salience linear mixed model, there was no difference in SCR AUC between stimulus modalities for intensity or unpleasantness, suggesting that these ratings also track with physiological arousal, as they are related to salience and are often correlated in healthy adults.

**Mediation Analysis**

Stimulus intensity predicted intensity ratings (*a* = 15.42, p < 2e^-16^), and, as shown earlier, stimulus intensity predicted SCR AUC (*c* = 0.44, p < 1.17e^-09^). When accounting for intensity as a mediator (Figure S2A), intensity ratings predicted SCR AUC (*b* = 0.023, p < 2e^-16^) and stimulus intensity no longer predicted SCR AUC (*c’* = 0.088, p = 0.25), indicating that the intensity rating is a complete mediator of this relationship, confirmed in the causal mediation analysis (*ab* = 0.35, p < 2e^-16^; proportion mediated= 80%). As shown in Figure S2B, stimulus intensity similarly predicted unpleasantness (*a* = 14.42, p < 2e^-16^), unpleasantness predicted SCR AUC (*b* = 0.023, p < 2e^-16^), and unpleasantness completely mediated the relationship between stimulus intensity and SCR AUC (*ab* = 0.33, p < 2e^-16^; proportion mediated = 76%), as the effect diminished when unpleasantness was accounted for in the mediation analysis (*c’* = 0.11, p = 0.16).
