## Supplementary material for "Subjective salience ratings are a reliable proxy for physiological measures of arousal": Figure S1

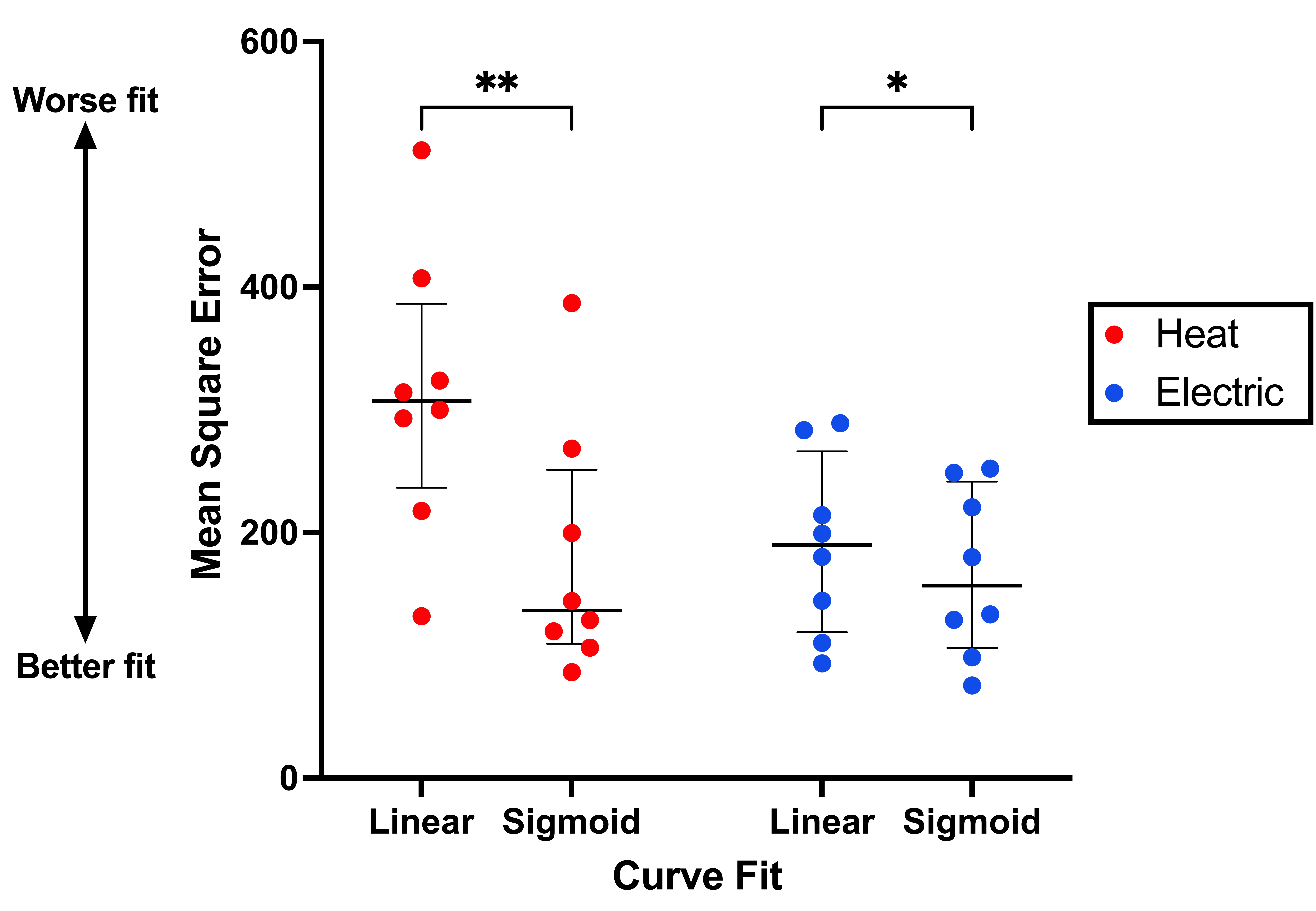
**Figure S1. Mean Squared Error of linear and sigmoid models compared using leave-one-out cross validation for electric (blue) and heat (red) stimuli.** Dots represent the MSE per regression type per participant across stimulus modality. Horizontal bars reflect the group median and error bars reflect the interquartile range.
