## Supplementary material for "Subjective salience ratings are a reliable proxy for physiological measures of arousal": Figure S2

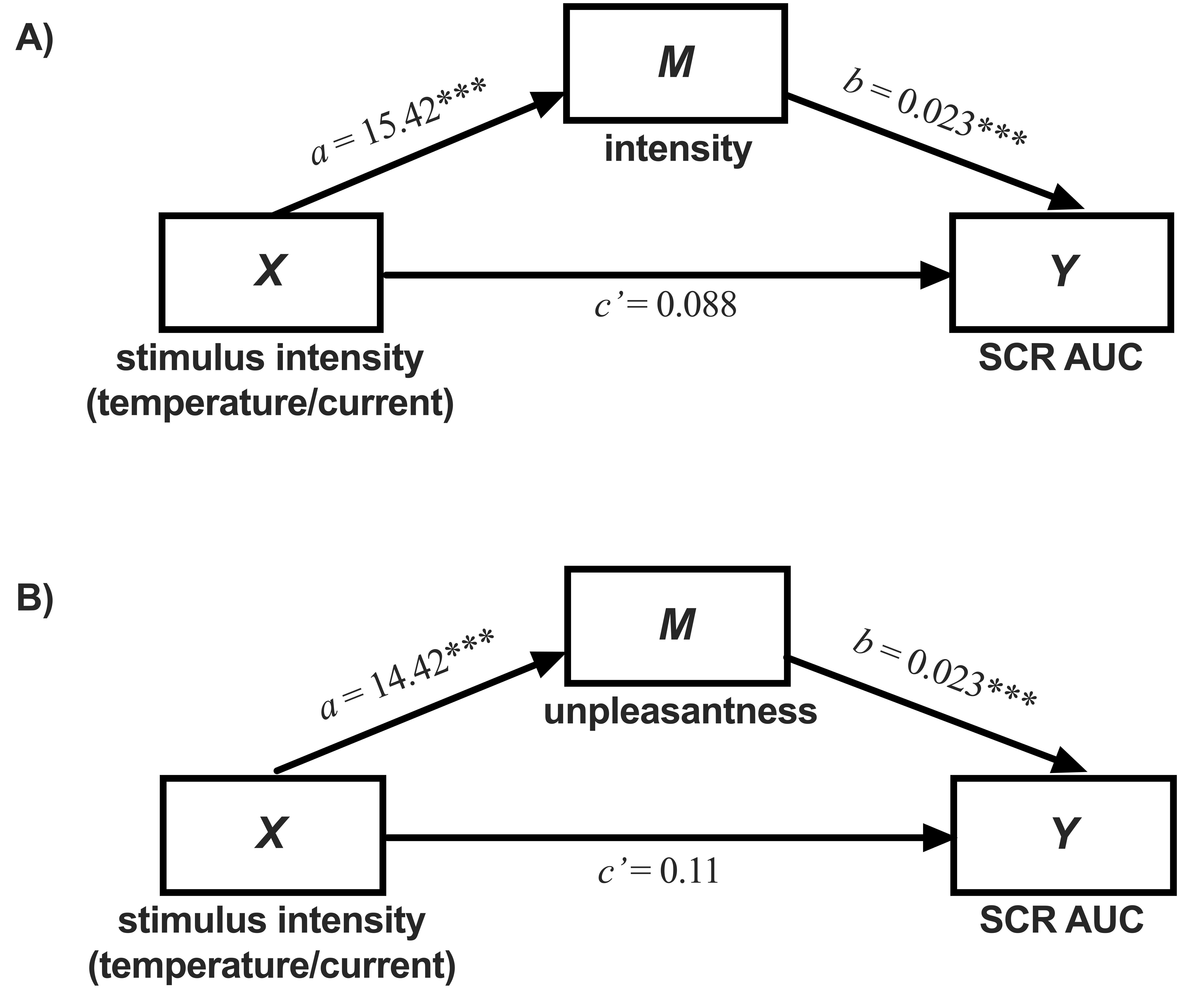

**Figure S2. (A) Intensity ratings and (B) unpleasantness ratings both completely mediated the relationship between stimulus intensity and SCR AUC.**
