## Supplementary material for "Subjective salience ratings are a reliable proxy for physiological measures of arousal": Table S1

**Table S1: Bayesian Repeated Measures ANOVA Model Comparison for Bayes Factor Determination Analysis for Sample Size.**

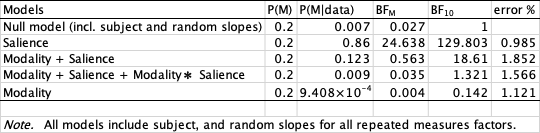
