## Supplementary figures and images for "Subjective salience ratings are a reliable proxy for physiological measures of arousal"

### Table S2

**Table S2: Analysis of Effects of Bayesian Repeated Measures ANOVA.**

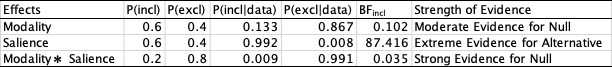
